# Bioinformatic-based identification of 3 immune-related genes for prognostic in breast invasive carcinoma

**DOI:** 10.1101/2022.09.23.509179

**Authors:** Xiaoying Cui, Shuang Song, Wenjuan Zhang, Dawei Wang, Zhijun Fang

**Author notes:** Address correspondence to Zhijun Fang, or Dawei Wang.

## Abstract

Breast invasive carcinoma (BRCA) is the leading cause of cancer deaths in female worldwide. Immune cell infiltration is considered to be crucial factor for the success or failure of BRCA therapy. As we all known, early diagnosis of breast cancer can greatly improve the survival rate of patients. Hence, to screen effective biomarkers for breast cancer immunotherapy might be extremely important. In this research, we identified an immune-related, three-gene biomarkers for prognosis of BRCA. We obtained altogether 192 differentially expressed genes (DEGs) from GEO datasets (GSE22820, GSE36295) and GEPIA database, followed by the Gene Ontology (GO) and Kyoto Encyclopedia of Genes and Genomes (KEGG) analysis. We screened 3 immune-related candidate biomarkers composing by CXCL2, CXCL9 and RBP7 that improves survival prediction outcome in BRCA. Kaplan–Meier analysis was conducted to analyze the patient survival based on 3 genes. In addition, we found that selected biomarkers were closely connected with infiltration levels of different tumor immune cells. Moreover, principal component analysis showed three biomarkers could effectively distinguish tumor samples from normal samples. In summary, these findings suggested that CXCL2, CXCL9 and RBP7 are viable prognostic and diagnostic biomarkers and provide new ideas for immunotherapy solutions of BRCA in the future.

## Introduction

According to the latest global cancer statistics released in 2020, nearly 2.3 million people are diagnosed with BRCA and 685,000 patients die. BRCA has surpassed lung cancer as the most common cancer for the first time(Sung et al. 2021). Although the level of diagnosis and treatment have been substantially improved, incidence rate is still increasing annually and the prognosis of some patients is still poor. Increasing evidences proved that pathogenesis and progression of female breast cancer were influenced by multiple genes and cellular pathways(Liu et al. 2019). For the purpose of discovering more effective diagnostic and therapeutic strategies, there is an urgent need for screening new sensitive biomarkers.

Breast malignant tumor originates from epithelial tissue. BRCA contains not only of neoplastic cells, but also involves tumor microenvironment (TME), such as alterations of immune cells, extracellular matrix and other matrix components(Soysal et al. 2015; Choi et al. 2018; Lim et al. 2018; Wu et al. 2019). In the past few years, immunotherapy has been verified as promising treatment solutions and was widely applied to cancer medical therapy(Gasser et al. 2017). The most attractive approach of immunotherapy care therapies in cancer is immune checkpoint inhibitors (ICIs)(Su et al. 2016; Pardoll 2012).In 2018, BRCA has officially entered the age of immunotherapy. Although breast cancer immunotherapy is still in its infancy, there has been a rapid increase in breast cancer immunotherapy drug trials in recent years(Emens 2012). However, the benefits of this new treatment are only limited to few cases. Therefore, it is necessary to make further efforts to determine the immune-related prognostic biomarkers and targets in BRCA for achieving precision immune treatment.

Microarray analysis technology was considered to be a powerful tool for screening key gene indicators to reveal the molecular mechanism of diseases. In the study, three gene expression profiles from open online database were used to explore and validate a three immune-related signature biomarkers for BRCA poor prognosis. Subsequently, We evaluated the role of CXCL2, CXCL9 and RBP7 in BRCA with gene expression comparison, pathological staging analysis, overall survival (OS), various immune cells infiltration and copy number variation (CNV) data. The study shed light on three-gene combination could become a novel diagnostic biomarkers for predicting BRCA patients’ clinical outcomes in BRCA immunology.

## Materials and Methods

### Data Source and Processing

The corresponding GEO datasets contain mRNA expression profile of BRCA and normal clinical samples was selected. The target microarray datasets, including GSE22820 and GSE36295, were extracted from the Gene Expression Omnibus (GEO) database (https://www.ncbi.nlm.nih.gov/geo/).The GSE22820 dataset based on the GPL6480 platform (Agilent-014850 Whole Human Genome Microarray 4×44K G4112F) contained 176 tumor and 10 normal tissues. The GSE36295 dataset based on the GPL6244 platform (HuGene-1_0-st] Affymetrix Human Gene 1.0 ST Array) contained 45 tumor and 5 normal tissues.

1560 differential expression genes and the top 500 survival related lists in BRCA were obtained from the Gene Expression Profiling Interactive Analysis (GEPIA) (http://gepia2.cancer-pku.cn/#index), which contained the mRNA-seq expression data of 9736/8587 tumor/normal samples from the Genotype-Tissue Expression (GTEx) project, as well as TCGA database(Tang et al. 2019; Tang et al. 2017).It provides various functions including tumor/normal differential expression analysis, pathological stage, survival analysis, etc. A list of immune related genes were downloaded from the ImmPort database (https://immport.niaid.nih.gov) (Bhattacharya et al. 2014), which provided a total of 1793 human genes.

### Screening of differentially Expressed Gene (DEGs)

The GEO2R web tool was applied to identify the DEGs of GSE22820 and GSE36295 with the criteria of adjusted *P*-value (Adj.*P*)< 0.05 and | log2 fold-change(FC) | > 1.5. Then, the differential expression genes list were downloaded from GEPIA with the same threshold values. The genes with log_2_ FC > 1.5 were defined as up-regulated genes and the genes with log_2_ FC < -1.5 were defined as down-regulated genes. We made a overlap and extract the differentially expressed genes using the Venn online web program (https://bioinfogp.cnb.csic.es/tools/venny/).

### Enrichment analysis of DEGs

The online Hiplot (https://hiplot.com.cn/) was used to perform GO (Ashburner et al. 2000) and KEGG(Kanehisa and Goto 2000) pathway analysis with the P< 0.05. The 10 most significant biological process (BP), cellular component (CC) along with molecular function (MF) terms for DEGs are displayed in the bubble charts. KEGG pathway analysis was used to reveal signaling pathways, which the DEGs might involve.

### Identification of DEGs related to immunity and prognosis

192 DEGs、survival related genes list from GEPIA and immune related genes list from ImmPort were conducted a Venn diagrams using the Venn online web program (https://bioinfogp.cnb.csic.es/tools/venny/). We found 3 DEGs related to immunity and prognosis in the final aggregated gene set.

### Validation of 3 target DEGs signature

Kaplan–Meier survival analysis was performed to evaluate the correlation between gene expression levels and the overall survival time in the BRCA patients by the use of GEPIA tool.

UALCAN (http://ualcan.path.uab.edu/analysis.html) is a comprehensive, user-friendly, interactive web resource using TCGA transcriptome data and clinical data for 31 cancer types. It was designed to provide the differential expression of tumor tissue and normal tissue, tumor stage, lymph node metastasis, and other related clinical parameters (Chandrashekar et al. 2017). We validated 3 target DEGs by using the UALCAN database, reanalyzed their expression differences in BRCA and normal tissue samples, and conducted correlation analysis between 3 target DEGs and age, race, and stage of lymph node metastasis. Multiple-gene comparison、 principal component analysis and gene expression among tumor stages for the three biomarker candidates were also conducted using the GEPIA.

### Relationship between 3 target DEGs with immune infiltration

Tumor Immune Estimation Resource(TIMER; https://cistrome.shinyapps.io/timer/) database is an effective web server for systematical analysis of immune infiltrates across diverse cancer types(Li et al. 2017). The latest version TIMER2 (http://timer.cistrome.org) was used to obtain correlation matrices in which tumor infiltration was calculated through many different methods, such as EPIC, CIBERSORT and XCELL (Li et al. 2020). The correlations of 3 target DEGs expression in BRCA with 6 immune cells ((B cells, CD4+ T cells, CD8+ T cells, neutrophils, macrophages, and dendritic cells (DCs)) were verified via the TIMER online approach. The correlation analysis was evaluated with the Spearman method. Furthermore, relationship between somatic copy number alterations(SCNA) and the abundance of immune infiltration were explored.

## Results

### Identification of 192 DEGs from three datasets

A workflow chart of the project design is presented in Figure 1. GSE22820 and GSE36295 available in the Gene Expression Omnibus (GEO) database together with differential expression genes available in the GEPIA database were subjected to overlapping analyses. The details of three expression datasets were shown in Table 1 and Figure 2A,2B. 192 commonly overlapping DEGs were identified totally(Figure 2C). After filtering out, 74 up-regulated and 118 down-regulated genes were discovered. Clearly, matrix metalloproteinase 11(MMP11) showed the most significant up-regulation level (log_2_ FC = 6.49), whereas leptin (LEP) showed the most significant down-regulation level (log_2_ FC = −5.48).

**Table 1.** Characteristics of datasets included in the identification of DEGs.

| Database ID | Platform ID | Number of Samples |  | Gene |
| --- | --- | --- | --- | --- |
|  |  | Tumor | Normal |  |
| GSE 22820 | GPL6480 | 176 | 10 | 1837 |
| GSE 36295 | GPL6244 | 45 | 5 | 337 |
| GEPIA | - | - | - | 1560 |

**Figure 1.**
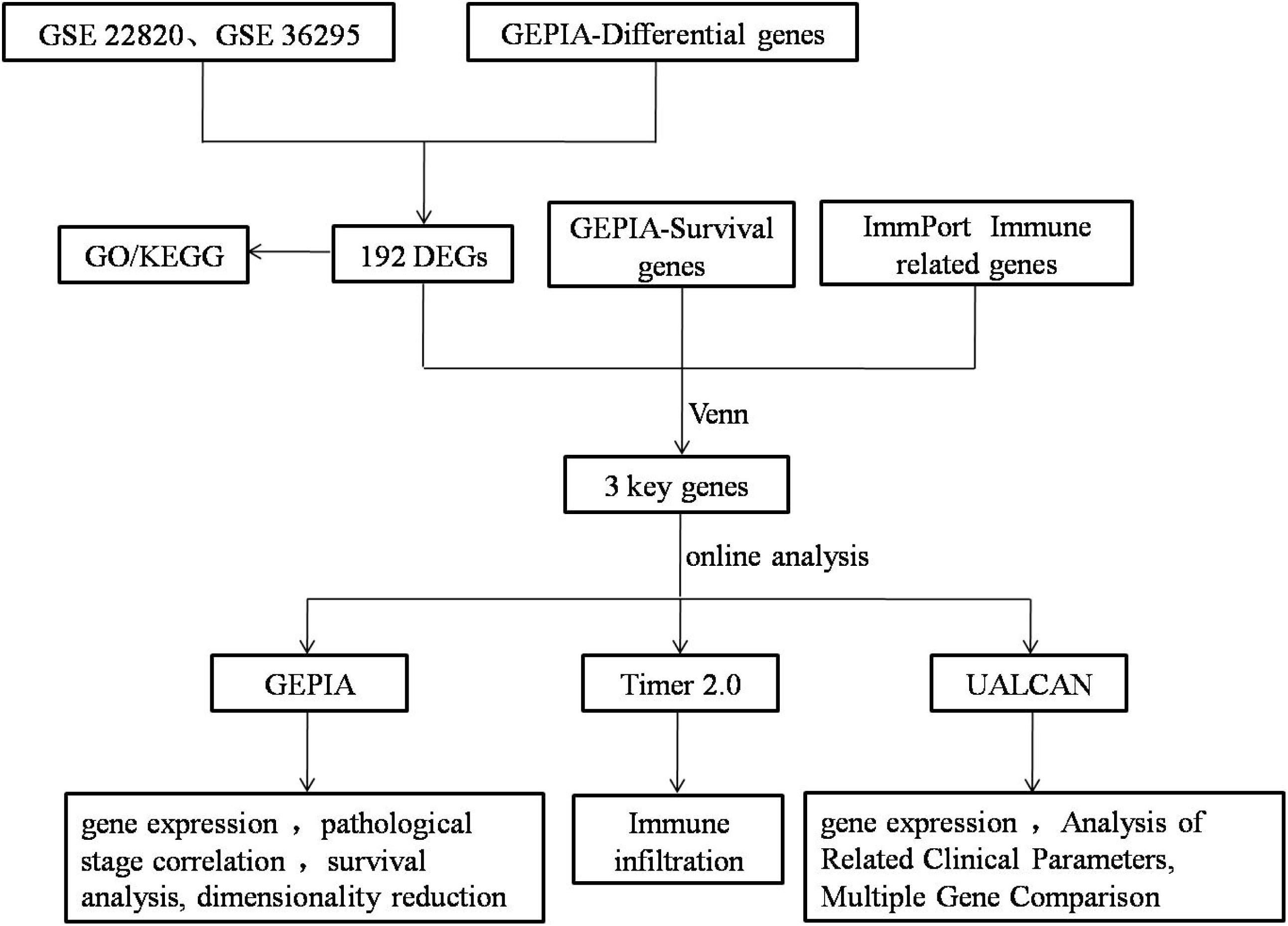
The work flow chart in this research project.

**Figure 2.**
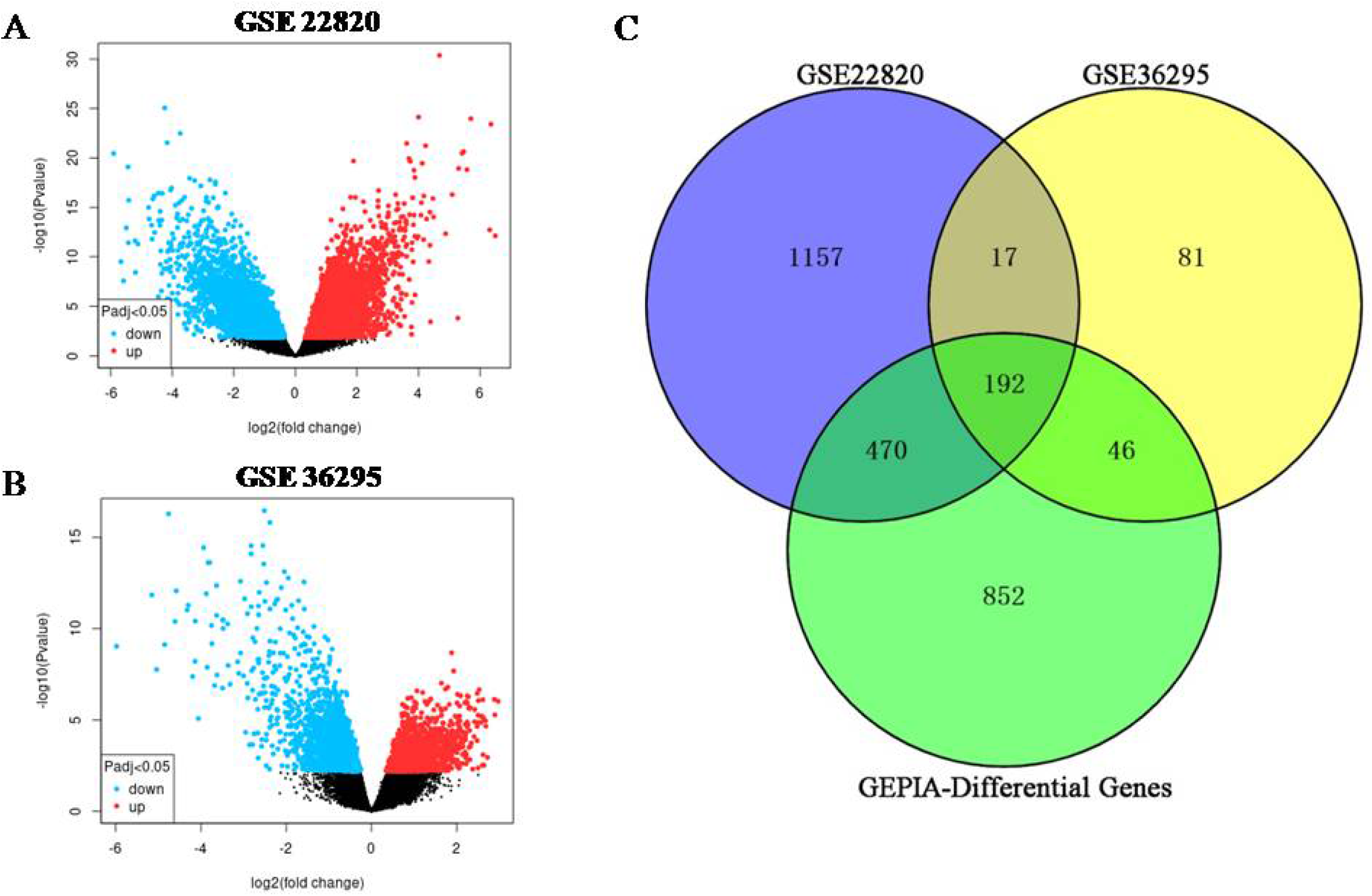
Identification of commonly DEGs in BRCA from public datasets. (A-B) Volcano plots of GSE22820 and GSE36295 datasets. (C) A total of 192 DEGs were identified and extracted from the three databases.

### Functional and Pathway Enrichment Analysis

We next imported the identified 192 DEGs into the Hiplot database to create a functional enrichment analysis. GO annotation contains three aspects of biological process (BP), cell composition (CC), and molecular function (MF). For BP category, nuclear division, organelle fission, mitotic nuclear division were the mainly significant terms (Figure 3A). For CC category, spindle, collagen-containing extracellular matrix, midbody and condensed chromosome were the mainly significant terms (Figure 3B). Besides, for MF category, glycosaminoglycan binding, microtubule binding, tubulin binding and sulfur compound binding were the mainly significant terms (Figure 3C).At the meanwhile, KEGG analysis showed that 192 DEGs are frequently enriched in multiple pathways consisted of PPAR signaling pathway, Cell cycle, AMPK signaling pathway, Progesterone-mediated oocyte maturation (Figure 3D).

**Figure 3.**
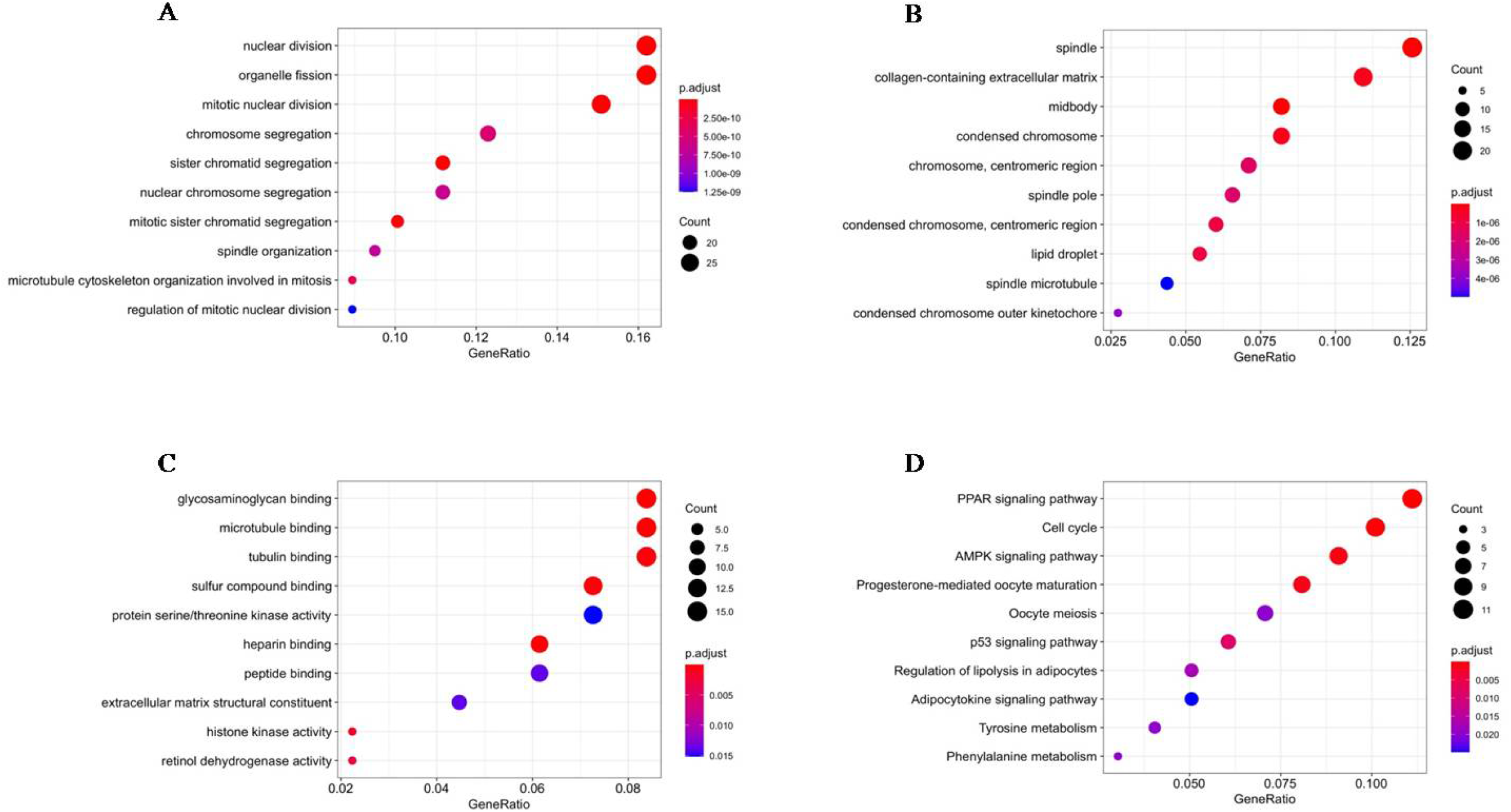
Gene enrichment analysis of integrated DEGs.(A) 10 significant enrichment results of BP. (B) 10 significant enrichment results of CC. (C) 10 significant enrichment results of MF. (D) 10 significant enrichment results of KEGG pathway.

### Screening of Immune Related DEGs with Prognostic Value

Under the criteria that adjusted P value is less than 0.05 and |log_2_ FC| more than 1.5, three immune genes related to BRCA prognosis were eventually verified as CXCL2, CXCL9, and RBP7,showed in the Venn diagram (Figure 4). The genetic information of three candidate biomarkers were demonstrated in Table 2.

**Table 2.** Basic characteristics of the 3 target genes.

| Gene | Description | Function | Location | Exon count |
| --- | --- | --- | --- | --- |
| CXCL2 | C-X-C motif chemokine ligand 2 | chemokine activity | 4q13.3 | 4 |
| CXCL9 | C-X-C motif chemokine ligand 9 | chemokine activity | 4q21.1 | 4 |
| RBP7 | retinol binding protein 7 | transporter activity | 1p36.22 | 5 |

**Figure 4.**
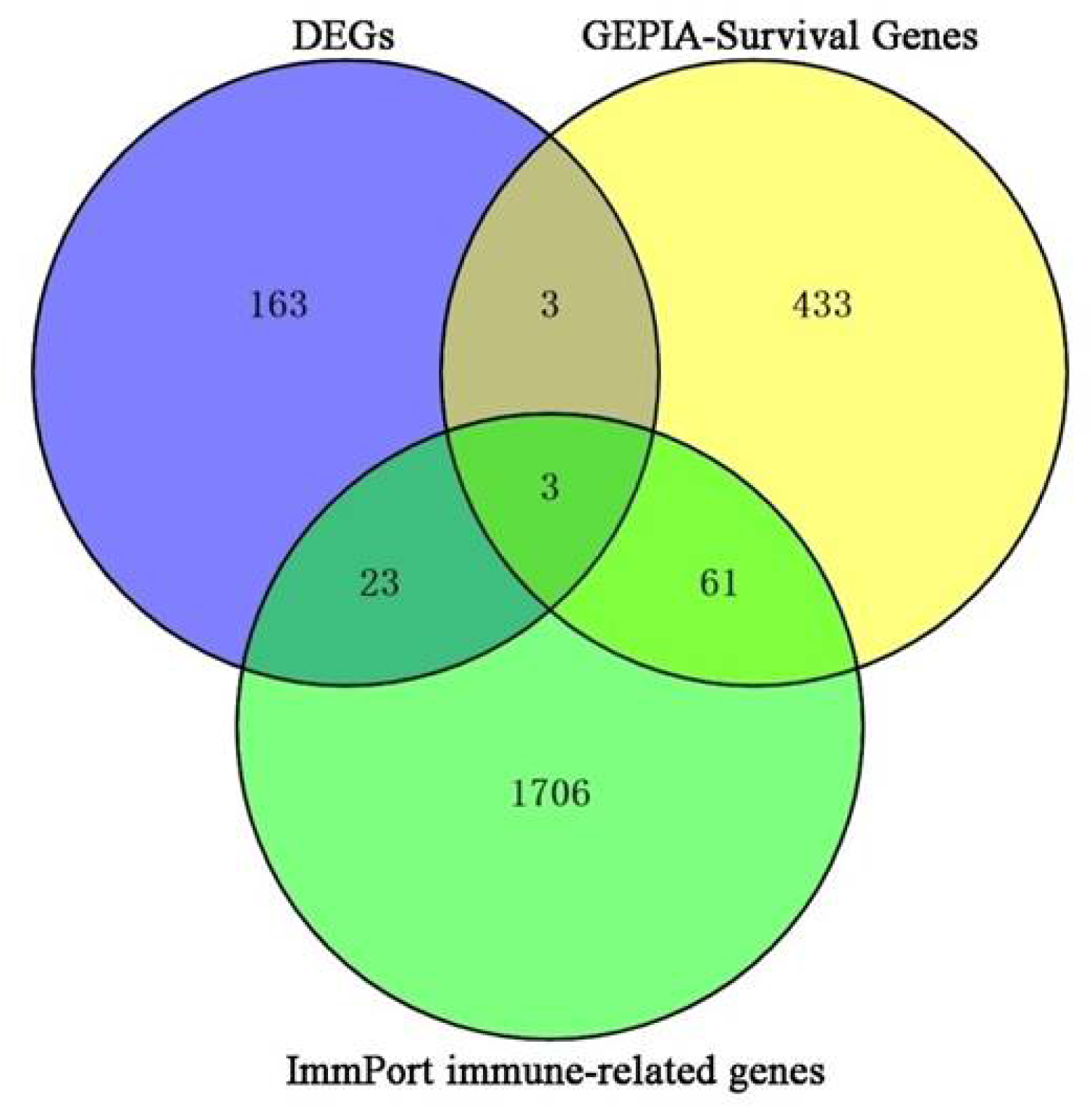
Intersection of DEGs, survival genes from GEPIA and immune genes from ImmPort.

### Validation of 3 Candidate Biomarkers

Using GEPIA, gene expression profiles of the 3 candidate biomarkers in BRCA tumor and normal specimens were exhibited in Figure 5A.The relative expression levels of CXCL2 and RBP7 in BRCA samples were significantly lower compared with non-BRCA samples, and CXCL9 expression in BRCA samples were obviously higher. The expression level of CXCL2 differed significantly in tumor stages (P < 0.05) (Figure 5B). In terms of survival analysis, We conducted survival analysis on the expression of 3 target DEGs and the overall survival time with hazard ratio (HR) and corresponding 95% confidence interval. The result revealed that CXCL9 up-regulation、CXCL2 and RBP7 down-regulation have a dismal effect on prognosis (Figure 5C).

**Figure 5.**
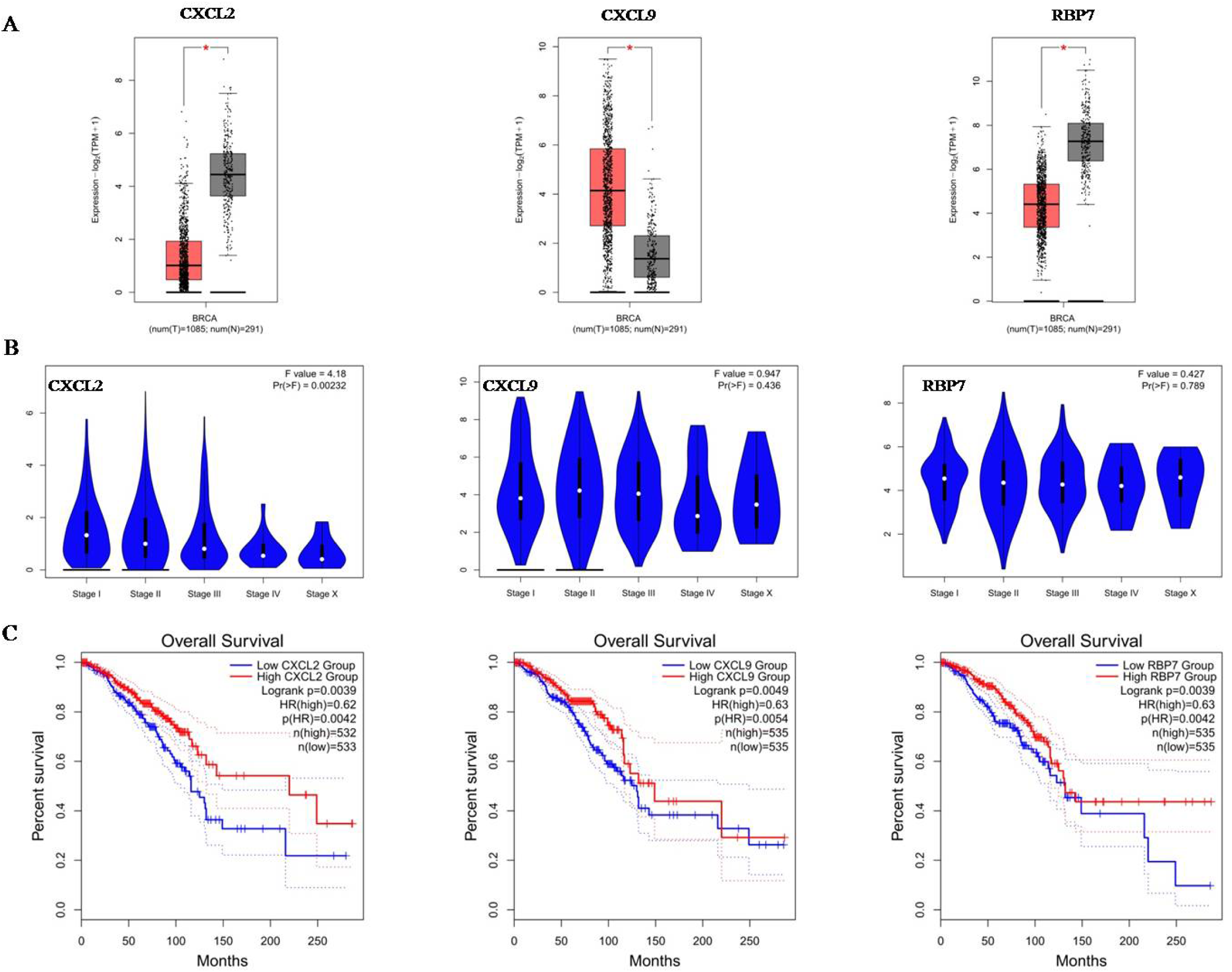
Prognostic biomarkers validation.(A) Differential expression between cancer and normal specimens. (B) Gene expression among tumor stages. (C) Overall survival curve.

### Association of gene expression with immune infiltration

In order to further explore the association between candidate biomarkers and the level of immune infiltration in BRCA, the TIMER was used according to the description of previous studies(Danaher et al. 2017; Siemers et al. 2017; Sousa and Maatta 2016). Noteworthy, all three genes were negatively correlated with tumor purity. It has been shown that the expression of three target biomarkers were all positively related to CD4+ T cells. The expression of CXCL9 was significantly related to immune infiltration and positively associated with all six types of immune infiltration(Figure 6A). In general, closely relationship was discovered between target genes expression and tumor immune infiltration. As shown in Figure 6B, the copy number variation (CNV) of CXCL2 and CXCL9 were correlated with infiltration of CD4+ T cells and macrophages. Meanwhile, RBP7 was directly correlated with CD8+ T cells, CD4+ T cells, macrophages, neutrophils, and DCs.

**Figure 6.**
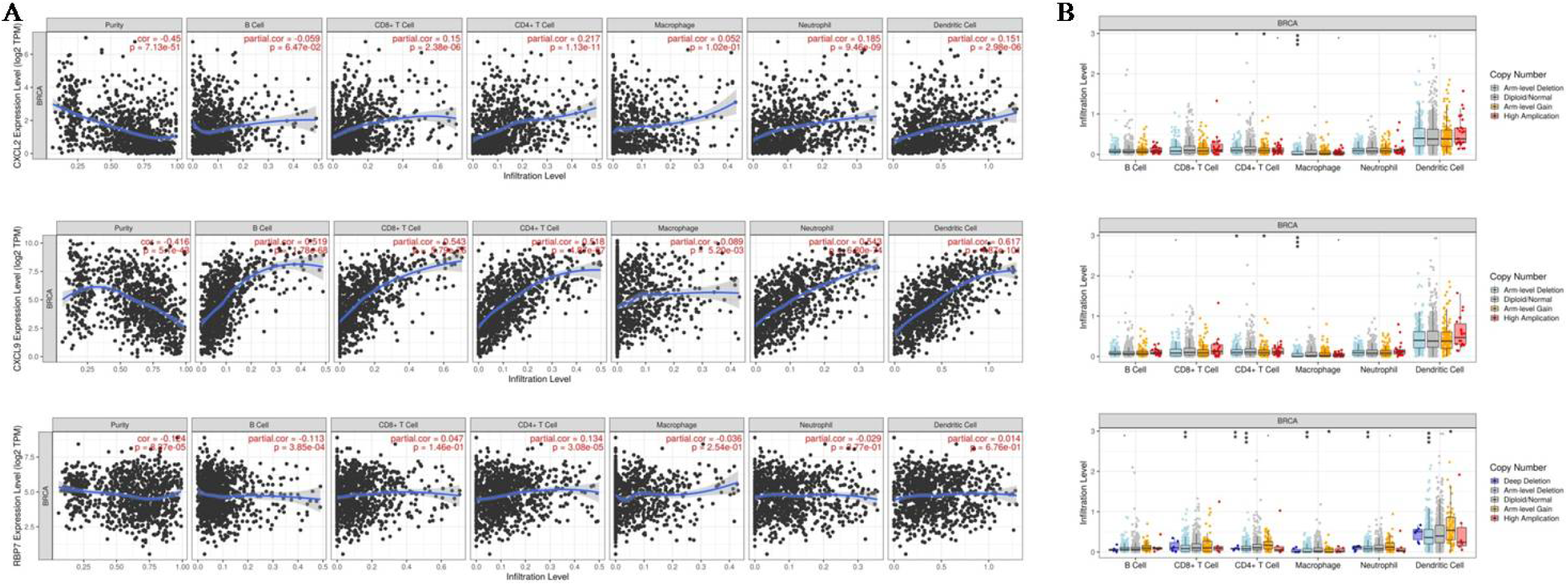
Association of biomarkers with immune cell infiltration. (A)The effect of CXCL2, CXCL9, and RBP7 on immune infiltration. (B) Relationship between SCNA and infiltration levels of immune cells.

### Clinical Analysis of CXCL2,CXCL9,RBP7

With UALCAN software, we detected the expression levels of CXCL2, CXCL9 and RBP7.There were obvious differences in the expression of these genes in the normal group and tumor group. CXCL9 was found to be highly expressed, while CXCL2 and RBP7 were significantly lower in tumor samples consistent with the bioinformatics results shown in Figure 5A (Figure 7).

**Figure 7.**
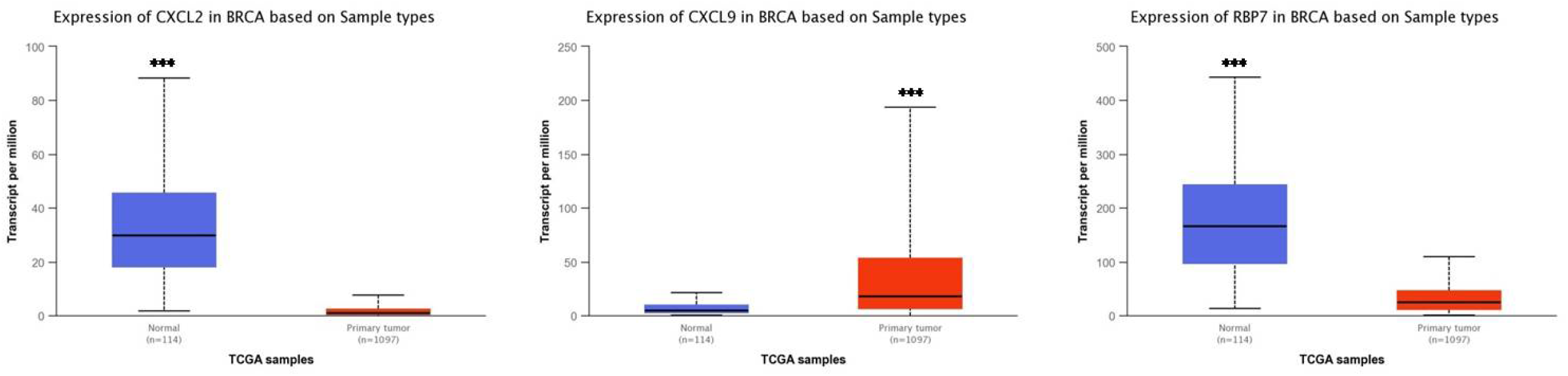
Comparison of expression of CXCL2, CXCL9, and RBP7 between tumor and normal tissues (***P < 0.001).

Subsequently, The influence of 3 genes expression on various clinicopathological characteristics including tumor stage, lymph node metastases, and age was investigated through UALCAN. The changes of transcript levels are consistent with analysis result we mentioned above. The expression levels of 3 genes all obviously increased when comparing tumor stages, lymph node metastasis, and age(Figure 8).

**Figure 8.**
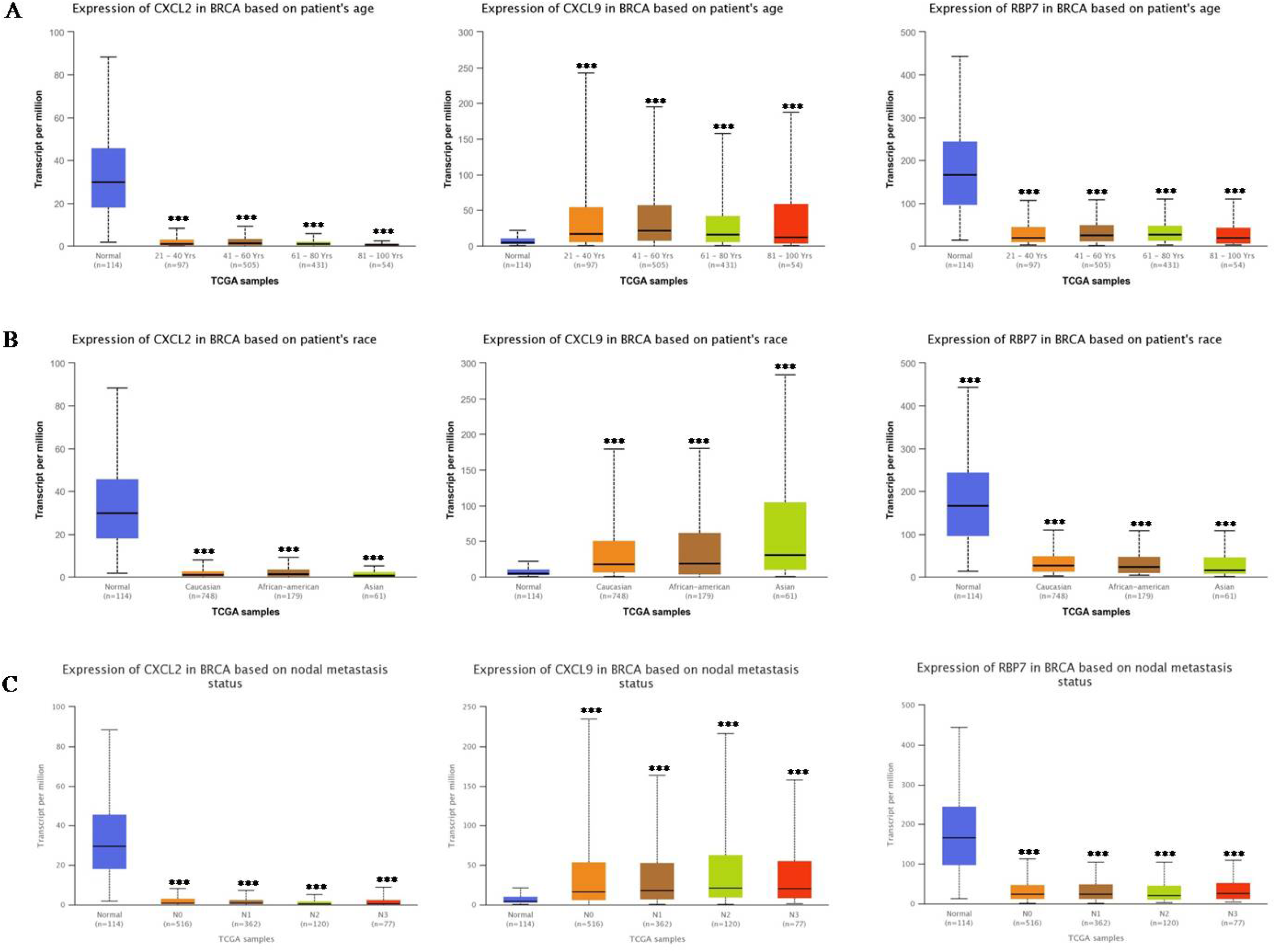
The correlation analysis between the gene relative expression of CXCL2, CXCL9, and RBP7 in BRCA and clinical parameters (***P < 0.001).

### Potential Diagnosis Biomarker for BRCA

Multiple-gene comparison analysis was evaluated by GEPIA. Among the three genes, the expression level of RBP7 was the highest, followed by CXCL9 and CXCL2(Figure 9).

**Figure 9.**
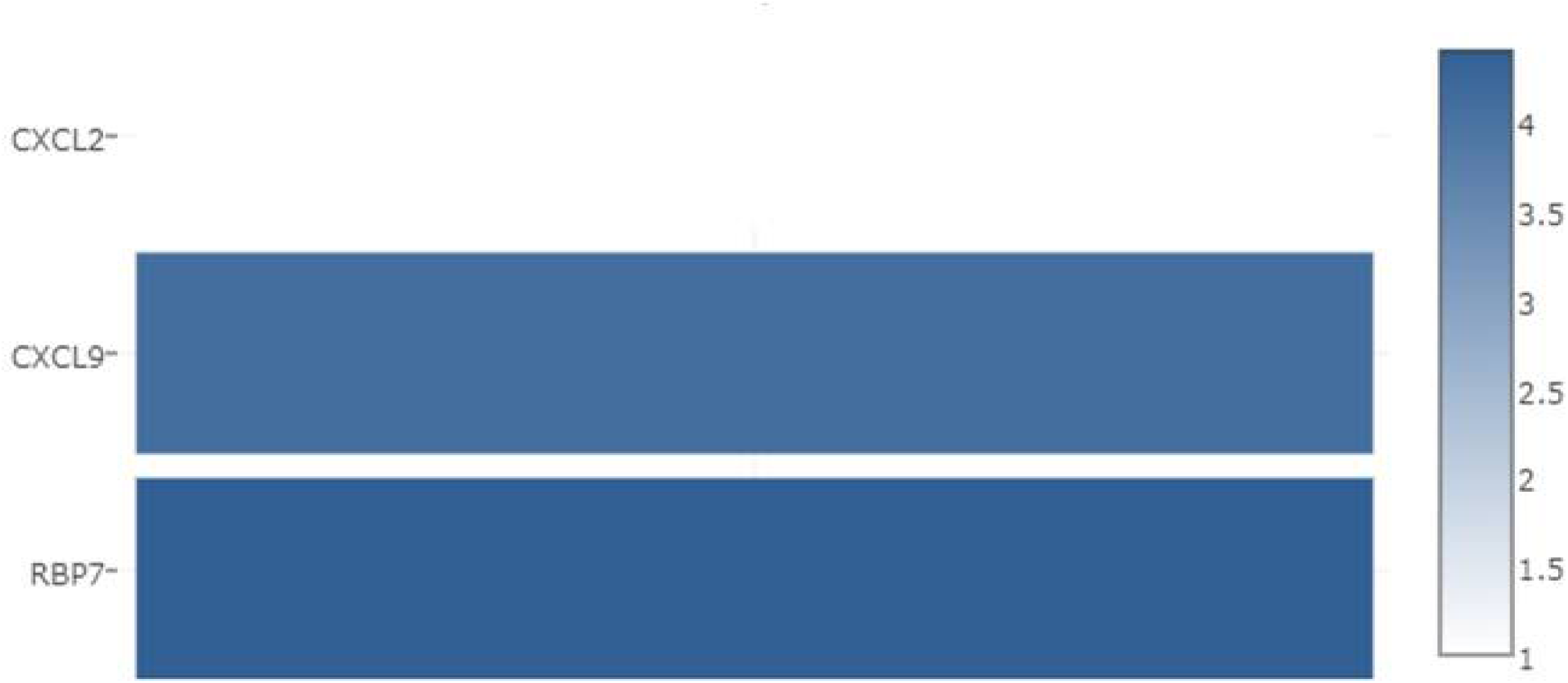
Multiple-gene comparison analysis for three candidate genes.

Principal component analysis of the three genes was conducted with TCGA tumor data, TCGA normal data, and GTEx data. In line with our expected result, combination of three genes could effectively distinguish between BRCA subjects and normal subjects (Figure 10), suggesting the potential possibility in diagnosis of BRCA.

**Figure 10.**
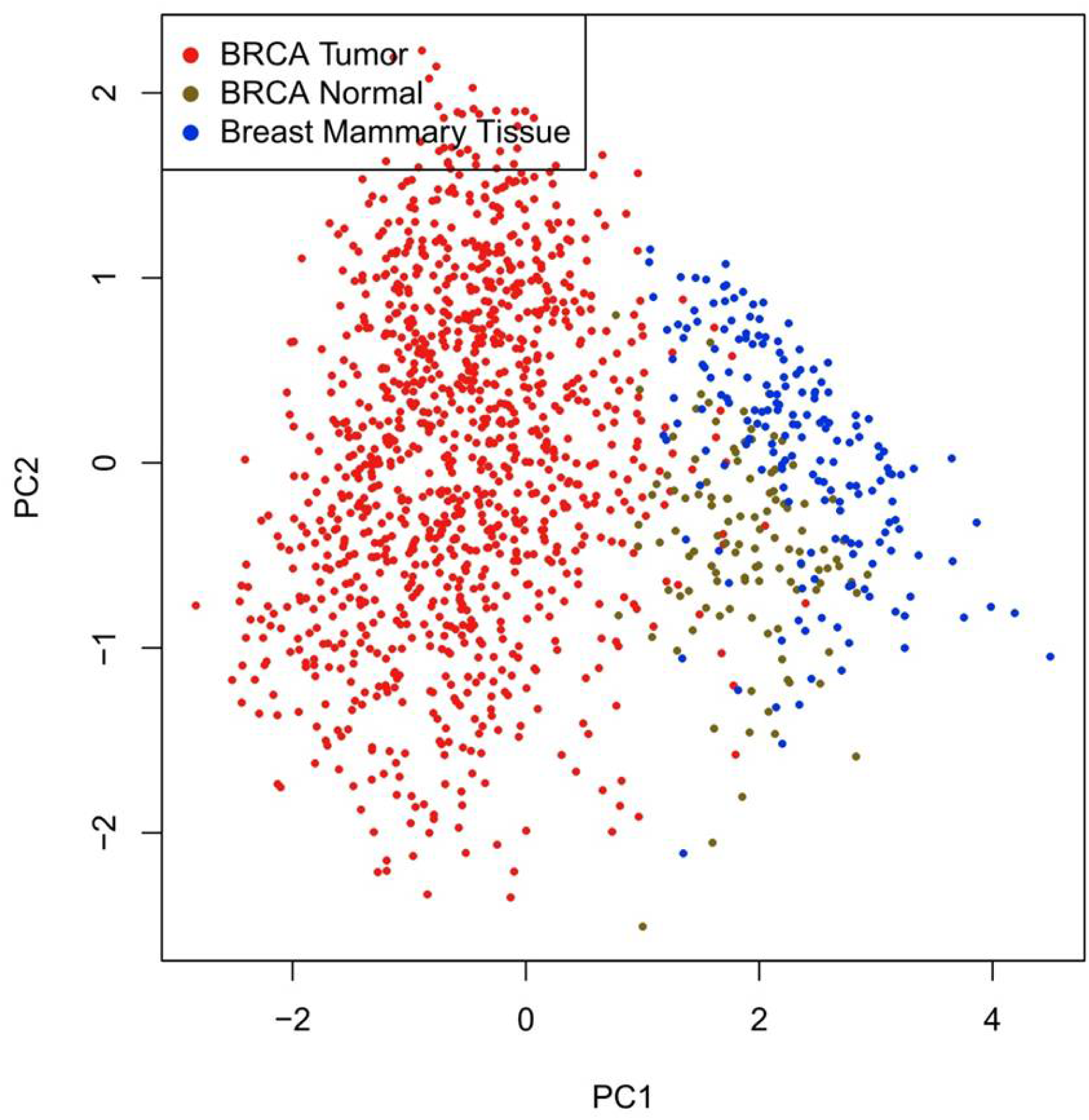
Principal component analysis of three candidate genes.

## Discussion

BRCA is the most common malignancy in women worldwide. Despite improvement in BRCA diagnosis and detection, a high recurrence rate still exist. Nearly 5–10% of breast cancer patients might have metastatic properties when it was found and 13% of cases with primary stage IV breast cancer survive ∼10 years(Eng et al. 2016). Therefore, early diagnosis of breast cancer is the key to improving the survival rate of patients. Based on the BRCA datasets from GEPIA and GEO database, we discovered 192 DEGs between BRCA and normal samples, of which, 74 was up-regulation genes and 118 was down-regulation genes. For GO terminology analysis, we found that 192 DEGs were mainly related to nuclear division and organelle. PPAR signaling pathway, cell cycle and AMPK signaling were the top three enriched pathways based on KEGG analysis. The above results shed lights on the molecular mechanism and treatment repertoires underlying BRCA.

The immune infiltration and tumor microenvironment have been reported to play pivotal roles in various cancers(Ju et al. 2020; Chen et al. 2019).There are many studies focus on identification of therapeutic targets and prognostic biomarkers, few biomarkers related to immune system were uncover in BRCA(Sharma et al. 2017; Zhu et al. 2019). CXCL2 CXCL9 and RBP7 were overlapped from the 192 DEGs, GEPIA and ImmPort online database. We found that these genes were markedly related to the prognosis of BRCA. All of them had a stronger association with the overall survival(OS) of breast tumor.

It is clear that chemokines and their receptors are involved in embryogenesis, hematopoiesis and malignant progression(Balkwill 2004). The chemokine (C-X-C motif) ligand (CXCL) family were recently reported to be involved in the development and progression of various cancer types (Cabrero-de Las Heras and Martinez-Balibrea 2018; Ding et al. 2016; Humblin and Kamphorst 2019; Lee et al. 2018; Zhang et al. 2017). Cytokines as well as their receptors were important players in the regulation of both induction and protection in breast cancer(Esquivel-Velazquez et al. 2015). Our research showed that CXCL2 had the strongest relationship with the pathological stages of breast cancer, while CXCL9 had weak association. An important mediators for leukocyte migration in inflammation, CXCL2 and RBP7 were observed to be down regulated in our research. Interestingly, previous studies had shown that CXCL2 was expressed at high levels in the bladder cancer tissues and ovarian cancer. We confirmed thatCXCL9 was over-expressed in BRCA patients compared with controls, consistent with previous studies(Narita et al. 2016).

The 134-amino acid protein, retinol-binding protein 7 (RBP7), a member of the fatty acid binding protein (FABP) family that binds retinol(Vogel et al. 2001). As an intracellular RBP, it is necessary for vitamin A stability and metabolism(Napoli 2017). Recent evidence suggests that cellular retinol-binding protein (CRBP) members might be involve in colon carcinogenesis and cancer stem cell maintenance(Berry et al. 2014; Karunanithi et al. 2017), yet its role in BRCA remains unknown. The bioinformatics analysis of BRCA found that RBP7 was a under-expressed gene. We show that low levels of RBP7 expression were obviously linked to poor clinical outcomes. The accurate results of three genes abnormal expression and biological function was ultimately to be confirmed by molecular biological experiments.

More recently, tumor immunotherapy that improve quality of life and delay disease progression has been rapidly developed, and it is found that the immune system has a dual effect in breast cancer, both promoting and preventing tumorigenesis(Delitto et al. 2016; Emens 2012). Furthermore, the immune cell infiltration including T cells, natural killer (NK) cells macrophages, and dendritic cell within tumor microenvironment is implicated in breast tumor development(Fridman et al. 2011; Mugler et al. 2007; Treilleux et al. 2004). Our findings showed the expression levels of CXCL2, CXCL9 and RBP7 had a significantly positive correlation with CD4+ T cells. CXCL9 was especially sensitive to immune infiltration than the other two genes. In agreement with *Liang et al*(Liang et al. 2021)who demonstrated that CXCL9 play a potential role in modulating the infiltration of immune cells. Taken together, three genes(CXCL2, CXCL9 and RBP7) has been proposed as an important player in BRCA oncogenesis and could be served as potential immune-related diagnostic and prognostic marker. These genes were closely linked to the overall survival time and nodal metastasis. BRCA samples and normal samples could be effectively distinguished by three gene biomarkers. These findings provides new insights into the BRCA immunobiology and revealed that CXCL2, CXCL9 and RBP7 were potential immune therapeutic targets.

## References

Ashburner, M., C.A. Ball, J.A. Blake, D. Botstein, H. Butler et al., 2000 Gene ontology: tool for the unification of biology. The Gene Ontology Consortium. Nat Genet 25 (1):25–29.

Balkwill, F., 2004 Cancer and the chemokine network. Nat Rev Cancer 4 (7):540–550.

Berry, D.C., L. Levi, and N. Noy, 2014 Holo-retinol-binding protein and its receptor STRA6 drive oncogenic transformation. Cancer Res 74 (21):6341–6351.

Bhattacharya, S., S. Andorf, L. Gomes, P. Dunn, H. Schaefer et al., 2014 ImmPort: disseminating data to the public for the future of immunology. Immunol Res 58 (2-3):234–239.

Cabrero-de Las Heras, S., and E. Martinez-Balibrea, 2018 CXC family of chemokines as prognostic or predictive biomarkers and possible drug targets in colorectal cancer. World J Gastroenterol 24 (42):4738–4749.

Chandrashekar, D.S., B. Bashel, S.A.H. Balasubramanya, C.J. Creighton, I. Ponce-Rodriguez et al., 2017 UALCAN: A Portal for Facilitating Tumor Subgroup Gene Expression and Survival Analyses. Neoplasia 19 (8):649–658.

Chen, B., J. Lai, D. Dai, R. Chen, X. Li et al., 2019 JAK1 as a prognostic marker and its correlation with immune infiltrates in breast cancer. Aging (Albany NY) 11 (23):11124–11135.

Choi, J., J. Gyamfi, H. Jang, and J.S. Koo, 2018 The role of tumor-associated macrophage in breast cancer biology. Histol Histopathol 33 (2):133–145.

Danaher, P., S. Warren, L. Dennis, L. D’Amico, A. White et al., 2017 Gene expression markers of Tumor Infiltrating Leukocytes. J Immunother Cancer 5:18.

Delitto, D., S.M. Wallet, and S.J. Hughes, 2016 Targeting tumor tolerance: A new hope for pancreatic cancer therapy? Pharmacol Ther 166:9–29.

Ding, Q., Y. Xia, S. Ding, P. Lu, L. Sun et al., 2016 Potential role of CXCL9 induced by endothelial cells/CD133+ liver cancer cells co-culture system in tumor transendothelial migration. Genes Cancer 7 (7-8):254–259.

Emens, L.A., 2012 Breast cancer immunobiology driving immunotherapy: vaccines and immune checkpoint blockade. Expert Rev Anticancer Ther 12 (12):1597–1611.

Eng, L.G., S. Dawood, V. Sopik, B. Haaland, P.S. Tan et al., 2016 Ten-year survival in women with primary stage IV breast cancer. Breast Cancer Res Treat 160 (1):145–152.

Esquivel-Velazquez, M., P. Ostoa-Saloma, M.I. Palacios-Arreola, K.E. Nava-Castro, J.I. Castro et al., 2015 The role of cytokines in breast cancer development and progression. J Interferon Cytokine Res 35 (1):1–16.

Fridman, W.H., J. Galon, F. Pages, E. Tartour, C. Sautes-Fridman et al., 2011 Prognostic and predictive impact of intra- and peritumoral immune infiltrates. Cancer Res 71 (17):5601–5605.

Gasser, S., L.H.K. Lim, and F.S.G. Cheung, 2017 The role of the tumour microenvironment in immunotherapy. Endocr Relat Cancer 24 (12):T283–T295.

Humblin, E., and A.O. Kamphorst, 2019 CXCR3-CXCL9: It’s All in the Tumor. Immunity 50 (6):1347–1349.

Ju, Q., X. Li, H. Zhang, S. Yan, Y. Li et al., 2020 NFE2L2 Is a Potential Prognostic Biomarker and Is Correlated with Immune Infiltration in Brain Lower Grade Glioma: A Pan-Cancer Analysis. Oxid Med Cell Longev 2020:3580719.

Kanehisa, M., and S. Goto, 2000 KEGG: kyoto encyclopedia of genes and genomes. Nucleic Acids Res 28 (1):27–30.

Karunanithi, S., L. Levi, J. DeVecchio, G. Karagkounis, O. Reizes et al., 2017 RBP4-STRA6 Pathway Drives Cancer Stem Cell Maintenance and Mediates High-Fat Diet-Induced Colon Carcinogenesis. Stem Cell Reports 9 (2):438–450.

Lee, N.H., M. Nikfarjam, and H. He, 2018 Functions of the CXC ligand family in the pancreatic tumor microenvironment. Pancreatology 18 (7):705–716.

Li, T., J. Fan, B. Wang, N. Traugh, Q. Chen et al., 2017 TIMER: A Web Server for Comprehensive Analysis of Tumor-Infiltrating Immune Cells. Cancer Res 77 (21):e108.#x2013;e110.

Li, T., J. Fu, Z. Zeng, D. Cohen, J. Li et al., 2020 TIMER2.0 for analysis of tumor-infiltrating immune cells. Nucleic Acids Res 48 (W1):W509–W514.

Liang, Y.K., Z.K. Deng, M.T. Chen, S.Q. Qiu, Y.S. Xiao et al., 2021 CXCL9 Is a Potential Biomarker of Immune Infiltration Associated With Favorable Prognosis in ER-Negative Breast Cancer. Front Oncol 11:710286.

Lim, B., W.A. Woodward, X. Wang, J.M. Reuben, and N.T. Ueno, 2018 Inflammatory breast cancer biology: the tumour microenvironment is key. Nat Rev Cancer 18 (8):485–499.

Liu, F., Y. Wu, Y. Mi, L. Gu, M. Sang et al., 2019 Identification of core genes and potential molecular mechanisms in breast cancer using bioinformatics analysis. Pathol Res Pract 215 (7):152436.

Mugler, K.C., M. Singh, B. Tringler, K.C. Torkko, W. Liu et al., 2007 B7-h4 expression in a range of breast pathology: correlation with tumor T-cell infiltration. Appl Immunohistochem Mol Morphol 15 (4):363–370.

Napoli, J.L., 2017 Cellular retinoid binding-proteins, CRBP, CRABP, FABP5: Effects on retinoid metabolism, function and related diseases. Pharmacol Ther 173:19–33.

Narita, D., E. Seclaman, A. Anghel, R. Ilina, N. Cireap et al., 2016 Altered levels of plasma chemokines in breast cancer and their association with clinical and pathological characteristics. Neoplasma 63 (1):141–149.

Pardoll, D.M., 2012 The blockade of immune checkpoints in cancer immunotherapy. Nat Rev Cancer 12 (4):252–264.

Sharma, P., S. Hu-Lieskovan, J.A. Wargo, and A. Ribas, 2017 Primary, Adaptive, and Acquired Resistance to Cancer Immunotherapy. Cell 168 (4):707–723.

Siemers, N.O., J.L. Holloway, H. Chang, S.D. Chasalow, P.B. Ross-MacDonald et al., 2017 Genome-wide association analysis identifies genetic correlates of immune infiltrates in solid tumors. PLoS One 12 (7):e0179726.

Sousa, S., and J. Maatta, 2016 The role of tumour-associated macrophages in bone metastasis. J Bone Oncol 5 (3):135–138.

Soysal, S.D., A. Tzankov, and S.E. Muenst, 2015 Role of the Tumor Microenvironment in Breast Cancer. Pathobiology 82 (3-4):142–152.

Su, M., C.X. Huang, and A.P. Dai, 2016 Immune Checkpoint Inhibitors: Therapeutic Tools for Breast Cancer. Asian Pac J Cancer Prev 17 (3):905–910.

Sung, H., J. Ferlay, and R.L. Siegel, 2021 Global Cancer Statistics 2020: GLOBOCAN Estimates of Incidence and Mortality Worldwide for 36 Cancers in 185 Countries. 71 (3):209–249.

Tang, Z., B. Kang, C. Li, T. Chen, and Z. Zhang, 2019 GEPIA2: an enhanced web server for large-scale expression profiling and interactive analysis. Nucleic Acids Res 47 (W1):W556–W560.

Tang, Z., C. Li, B. Kang, G. Gao, C. Li et al., 2017 GEPIA: a web server for cancer and normal gene expression profiling and interactive analyses. Nucleic Acids Res 45 (W1):W98–W102.

Treilleux, I., J.Y. Blay, N. Bendriss-Vermare, I. Ray-Coquard, T. Bachelot et al., 2004 Dendritic cell infiltration and prognosis of early stage breast cancer. Clin Cancer Res 10 (22):7466–7474.

Vogel, S., C.L. Mendelsohn, J.R. Mertz, R. Piantedosi, C. Waldburger et al., 2001 Characterization of a new member of the fatty acid-binding protein family that binds all-trans-retinol. J Biol Chem 276 (2):1353–1360.

Wu, Q., J. Li, Z. Li, S. Sun, S. Zhu et al., 2019 Exosomes from the tumour-adipocyte interplay stimulate beige/brown differentiation and reprogram metabolism in stromal adipocytes to promote tumour progression. J Exp Clin Cancer Res 38 (1):223.

Zhang, H., Y.L. Ye, M.X. Li, S.B. Ye, W.R. Huang et al., 2017 CXCL2/MIF-CXCR2 signaling promotes the recruitment of myeloid-derived suppressor cells and is correlated with prognosis in bladder cancer. Oncogene 36 (15):2095–2104.

Zhu, J., P.F. Petit, and B.J. Van den Eynde, 2019 Apoptosis of tumor-infiltrating T lymphocytes: a new immune checkpoint mechanism. Cancer Immunol Immunother 68 (5):835–847.

